# The interplay of synaptic plasticity and scaling enables self-organized formation and allocation of multiple memory representations

**DOI:** 10.1101/260950

**Authors:** Johannes Maria Auth, Timo Nachstedt, Christian Tetzlaff

## Abstract

It is commonly assumed that memories about experienced stimuli are represented by groups of highly interconnected neurons called cell assemblies. This requires allocating and storing information in the neural circuitry, which happens through synaptic weight adaptation. It remains, however, largely unknown how memory allocation and storage can be achieved and coordinated to allow for a faithful representation of *multiple* memories without disruptive interference between them. In this theoretical study, we show that the interplay between conventional synaptic plasticity and homeostatic synaptic scaling organizes synaptic weight adaptations such that a new stimulus forms a new memory and where different stimuli are assigned to distinct cell assemblies. The resulting dynamics can reproduce experimental in-vivo data, focusing on how diverse factors as neuronal excitability and network connectivity, influence memory formation. Thus, the here presented model suggests that a few fundamental synaptic mechanisms may suffice to implement memory allocation and storage in neural circuitry.

## Introduction

Learning and memorizing information about the environment over long time scales is a vital function of neural circuits of living beings. For this, different elements of a neural circuit - the neurons and synapses - have to coordinate themselves to accurately form, organize, and allocate internal long-term representations of the different pieces of information received. While many processes and dynamics of the single elements are well documented, their coordination remains obscure. How do neurons and synapses self-organize to form functional, stable, and distinguishable memory representations? Moreover, what mechanisms underlie the self-organized coordination yielding such representations?

Current hypotheses centre on long-term synaptic plasticity as the core mechanism for robust coordination of the circuits’ elements into memory representations^1–4^. Different from this, in this theoretical study, we identify that formation of functional memory representations requires the interaction of synaptic plasticity with (slower) synaptic scaling, which is a mechanisms that enforces network homeostasis^5–7^. This interaction ensures the reliable, self-organized coordination of synaptic and neuronal elements into memories.

As discussed in the following, the organization of memory representations is subdivided into two processes: memory formation and allocation.

The reliable *formation* of memory, hence of internal stimulus representations in neural circuits, is explained by the Hebbian hypothesis. In brief, the Hebbian hypothesis^3, 8–11^ states that, when a neural circuit receives a new piece of information, the corresponding stimulus activates a group of neurons and, via activity-dependent long-term synaptic plasticity^1, 12–14^, adapts the weights or efficacies of synapses between these neurons. This adaptation remodels the activated group of neurons to a strongly interconnected group of neurons called Hebbian cell assembly (CA). This newly formed CA serves as an internal long-term representation (long-term memory) of the corresponding stimulus^3, 10, 11^. Recall of this memory translates into the activation of the respective CA. In order to recognize similar pieces of information, similar stimuli also have to be able to activate the corresponding CA. This is enabled by the strong recurrent interconnections between CA-neurons resulting in pattern completion^3, 15^. Several experimental and theoretical studies have investigated the formation and recall of CAs given synaptic plasticity^11, 16–22^. Recent theoretical studies already indicate that, in addition to synaptic plasticity, homeostatic mechanisms, such as synaptic scaling, are required to keep the neural circuit in a functional regime^23–26^. However, all the above-mentioned studies investigate the coordination of synaptic and neuronal dynamics within the group of neurons (memory formation), but they do not consider the dynamics determining which neurons are recruited to form this group, or rather why is the stimulus allocated to this specific group of neurons and not to others (memory allocation;^27^).

For a proper *allocation* of stimuli to their internal representations, the synapses from the neurons encoding the stimulus to the neurons encoding the internal representation have to be adjusted accordingly. Considering only these types of synapses ("feed-forward synapses"), several studies show that long-term synaptic plasticity yields the required adjustments of synaptic weights^28–33^ such that each stimulus is mapped to its corresponding group of neurons. However, theoretical studies^34, 35^, which investigate the formation of input-maps in the visual cortex^36, 37^, show that the stable mapping or allocation of stimuli onto a neural circuit requires, in addition to synaptic plasticity, homeostatic mechanisms such as synaptic scaling. Note that all these studies focus on the allocation of stimuli to certain groups of neurons; however, they do not consider the dynamics of the synapses within these groups ("recurrent synapses").

Thus, up to now, it remains unclear how a neural circuit coordinates in a self-organized manner the synaptic and neuronal dynamics underlying the reliable allocation of stimuli to neurons with the simultaneous dynamics underlying the proper formation of memory representations. If these two memory processes are not tightly coordinated, the neural system could show awkward, undesired dynamics: On the one hand, memory allocation could map a stimulus to a group of unconnected neurons, which impedes the formation of a proper CA. On the other hand, the formation of a CA could bias the dynamics of allocation such that multiple stimuli are mapped onto the same CA disrupting the ability to discriminate between these stimuli.

In this theoretical study, we show in a network model that long-term synaptic plasticity^10, 38, 39^ together with the slower, homeostatic processes of synaptic scaling^5–7, 40^ leads to the self-organized coordination of synaptic weight changes at feed-forward and recurrent synapses. Remarkably, the synaptic changes of the recurrent synapses yields the reliable formation of CAs. In parallel, the synaptic changes of the feed-forward synapses links the newly formed CA with the corresponding stimulus-transmitting neurons without interrupting already learned ones assuring the proper allocation of CAs. The model reproduces *in-vivo* experimental data and provides testable predictions. Furthermore, the analysis of a population model, capturing the main features of the network dynamics, allows us to determine three generic properties of the interaction between synaptic plasticity and scaling, which enable the formation and allocation of memory representations in a reliable manner. These properties of synaptic adaptation are that (i) synaptic weights between two neurons with highly-correlated activities are strengthened (homosynaptic potentiation), (ii) synaptic weights between two neurons with weakly-correlated activities are lowered (heterosynaptic depression), and (iii) the time scale of synaptic weight changes are regulated by the post-synaptic activity level.

## Results

Throughout this study, we consider a neural system receiving environmental stimuli (e.g., geometrical shapes as a blue triangle, S1, or a red square, S2), which evoke certain activity patterns within the input area (e.g., blue pattern I1 evoked by S1; 1 A). Each activity pattern, in turn, triggers via several random feed-forward synapses the activation of a subset of neurons in the recurrently connected memory area. For simplicity, all neurons of the system are considered to be excitatory; only a single all-to-all connected inhibitory unit (resembling an inhibitory population) in the memory area regulates its global activity level (1 B). Furthermore, as we investigate here neuronal and synaptic dynamics happening on long time scales, we neglect the influence of single spikes and, thus, directly consider the dynamics of the neuronal firing rates (see *Materials and Methods*). All synapses between the excitatory neurons (feed-forward and recurrent) are adapted by activity-dependent long-term plasticity. Due to the recurrent connections, the memory area should robustly form internal representations of the environmental stimuli, while simultaneously the feed-forward synapses should provide a proper allocation of the stimuli (activity patterns in input area) onto the corresponding representations. In the following, we will show in a step-by-step approach that the interplay of long-term synaptic plasticity (here conventional Hebbian synaptic plasticity) with homeostatic synaptic scaling (see Methods;^7, 23, 41^) coordinates synaptic changes such that proper formation and allocation of memory is ensured.

First, stimulus S1 is repetitively presented ten times (first learning phase; 1 C). Given this stimulation, the resulting synaptic adaptations of feed-forward and recurrent synapses should yield the proper formation of an internal representation indicating that the dynamics underlying memory allocation (changes of feed-forward synapses) does not impede the recurrent dynamics of CA-formation. After the formation of this representation, next, we repetitively present a different stimulus S2 (second learning phase). Due to this stimulation, the neural system should form another CA representing S2, which is independent of the first one. The proper formation of a second CA indicates that memory allocation is not biased by recurrent dynamics enabling a reliable discrimination between stimuli. Please note that we consider three test phases (1 C), during which synaptic dynamics are fixed, to enable the investigation of the resulting response dynamics of the circuit according to the different stimuli. Otherwise, the system is always plastic. In general, we expect that the neural system should form strongly interconnected groups of neurons according to each learning stimulus (memory formation; CA1 and CA2 in 1 D indicated by thicker lines) while remaining neurons in the memory area last weakly interconnected (RR). In addition, the synapses from the neurons in the input area, which are activated by a specific stimulus (I1 and I2), to the corresponding CAs should have larger weights while all other feed-forward connections remain rather weak (memory allocation; I1 to CA1 and I2 to CA2). After showing that the interplay of synaptic plasticity and scaling yields such a synaptic structure, in the next step, we derive a population model of the neural system and analyse the underlying synaptic and neuronal dynamics to identify the required generic properties determining the synaptic adaptations. Finally, we demonstrate that our theoretical model matches and provides potential explanations for a series of experiments revealing the relation between neuronal dynamics and the allocation of memory^42^. In addition, we compile some experimentally verifiable predictions, which would confirm the here-proposed hypothesis that the interplay between synaptic plasticity and scaling is required for the proper formation and allocation of memories.

**Figure 1.**
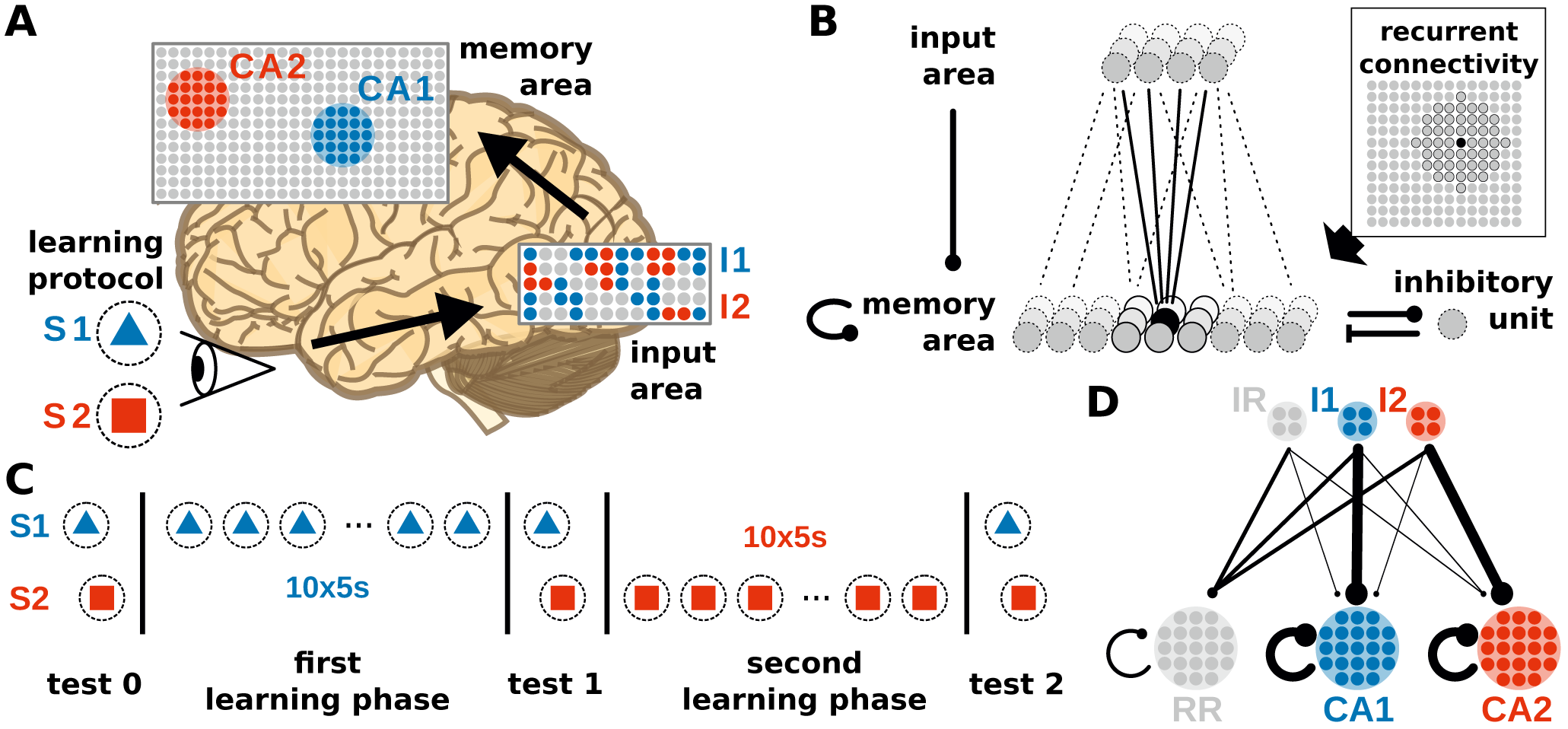
The neural system to investigate the coordination of synaptic and neuronal dynamics underlying the allocation and formation of memories consists of two areas receiving external stimuli. (A): The neural system receives different external stimuli (e.g., a blue triangle, S1, or a red square, S2) yielding the activation of subsets of neurons (colored dots) in the input (I1 and I2) and memory area (CA1 and CA2). (brain image by Hugh Guiney under license CC BY-SA 3.0) (B): Each excitatory neuron in the memory area receives inputs from a random subset of excitatory neurons being in the input area, from its neighbouring neurons of the memory area (indicated by the dark Gray units in the inset) and from a global inhibitory unit. All synapses between excitatory neurons are plastic regarding the interplay of long-term synaptic plasticity and synaptic scaling. (C): Throughout this study, we consider two learning phases during each a specific stimulus is repetitively presented. In addition, test phases are considered with stopped synaptic dynamics for analyses. (D): Schematic illustration of the average synaptic structure ensuring a proper function of the neural system. This structure should result from the neuronal and synaptic dynamics in conjunction with the stimulation protocol. IR and RR represent populations of remaining neurons being not directly related to the dynamics of memory formation and allocation. Details see main text.

### Formation and allocation of one memory representation

Before learning, feed-forward as well as recurrent synapses, on average, do not show any structural bias (2 A, test 0) such that the presentation of an environmental stimulus (e.g., S1 or S2) triggers via the activation of a stimulus-specific pattern within the input area (I1 or rather I2) the activation of a random pattern of active neurons in the memory area. To evaluate whether these activated neurons are directly connected with each other, which serves as the basis for the formation of a CA^3, 10^, we consider the average shortest path length (ASPL; Supplementary Section A) between these neurons. In general, the ASPL is a graph theoretical measure to assess the number of units (here neurons) along the shortest path to be taken in the network to reach one specific unit starting from another unit averaged across all pairs of units (here we consider only the pairs of activated neurons). Thus, as the ASPL-value is high, we conclude that the activated neurons in the memory area are not directly connected with each other (2 B, test 0); thus, the activity pattern is mainly determined by the random feed-forward connections.

By contrast, if a stimulus (here stimulus S1) is repeatedly presented in a given time interval, the neuronal and synaptic dynamics of the network reshapes the pattern of activated neurons in the memory area such that the final pattern consists of a group of interconnected neurons (decrease in ASPL; 2 B, test 1). As shown in our previous studies^23, 41^, the combination of synaptic plasticity and scaling together with a repeated activation of an interconnected group of neurons yields an average strengthening of the interconnecting recurrent synapses without significantly altering other synapses (2 A, test 1, bottom; neurons are sorted into groups retroactively; see Supplementary Figure S3 for exemplary, complete weight matrices). Taken together with the decreased ASPL this indicates the stimulus-dependent formation of a CA during the first learning phase. However, do the self-organizing dynamics also link specifically the stimulus-neurons with the CA-neurons? A repeated presentation of stimulus S1 yields an on average strengthening of synapses projecting from the stimulus-neurons I1 to the whole memory area (2 A, test 1, top). Essentially, the synapses from stimulus-I1-neurons to CA1-neurons have a significantly stronger increase in synaptic weights than the controls (CA2 and RR). In more detail, presenting stimulus S1 initially activates an increasing number of mainly unconnected neurons in the memory area (3; first presentation, fifth row, dark red dots; each dot represents the firing rate of one neuron in the memory area. Note that nearby neurons are connected with each other as indicated in the inset of 1 B and Supplementary Figure S1. This does not indicate the physical distance between neurons.). However, the ongoing neuronal activation during following stimulus presentations increases the recurrent synaptic weights (3, fourth row; each dot represents the average incoming weight of recurrent synapses a corresponding neuron receives) and also feed-forward weights from stimulus I1 to the neurons (3, second row; each dot represents the average incoming weight a neuron in the memory area receives from all feed-forward synapses from I1-neurons of the input area), which, in turn, increases the activity. This positive feedback loop between synaptic weights and neuronal activity leads to the emergence of groups of activated neurons, which are directly connected with each other (second presentation, fifth row). Such an interconnected group grows out, incorporating more directly connected neurons, until inhibition limits its growth (see below) and, furthermore, suppresses sparse activation in the remaining neurons by competition (fourth to tenth presentation). The recurrent weights among CA-neurons are increased until an equilibrium between Hebbian synaptic plasticity and synaptic scaling is reached (fourth row). Please note that, as our previous studies show^7, 23^, the interplay between these two mechanisms yields the existence of this equilibrium; otherwise synaptic weights would grow unbounded even if the neural activity is limited (see also detailed analysis below). Interestingly, the weights of the feed-forward synapses show a different dynamics as of the recurrent synapses. The average over all synaptic weights linking from the input area to each neuron in the memory area (first row) indicates that synapses connected to CA-neurons have a similar average weight compared to synapses connected to other neurons. This implies that the synaptic weight changes of the feed-forward connections to the CA-neurons are on average not significantly different than controls. However, if the feed-forward synapses are sorted according to the stimulus-affiliation of the pre-synaptic neuron, we see that only the weights of synapses from the S1-stimulus-neurons to the emerging CA-neurons are strengthened (second row, dark blue spot; see also 2 A, test 1, I1 to CA1), while weights of synapses from other stimulus-neurons to the CA-neurons are on average decreased (third row, white spot; see also 2 A, test 1, I2 to CA1). This implies a proper assignment of the stimulus to the newly formed CA during the learning phase resulting in a higher chance of activating the CA-neurons when the same stimulus is presented later again. These results reveal that the interaction of synaptic plasticity and scaling self-organizes for a wide parameter regime (see Supplementary Figure S2) synaptic changes at recurrent and feed-forward connections to form and allocate a memory representation in a previously random neuronal network.

**Figure 2.**
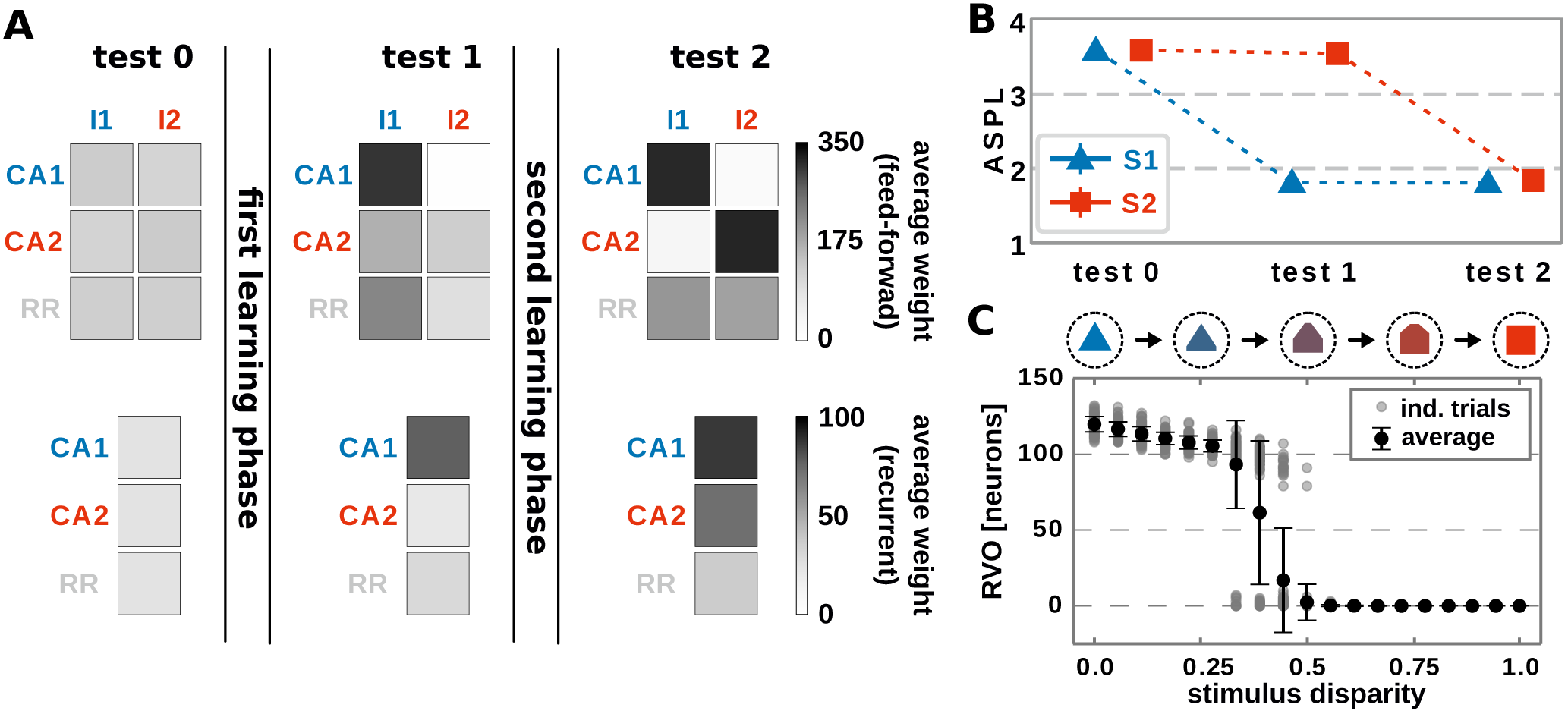
The interaction of conventional Hebbian synaptic plasticity and synaptic scaling enables the stimulus-dependent formation and allocation of memory representations in a neuronal network model. (A): During the test phases, the resulting network structure is evaluated. Top row: average synaptic weight of feed-forward synapses from input populations I1 and I2 activated by the corresponding stimuli to the groups of neurons which become a CA (CA1 and CA2) and others (RR). Please note that we first train the network, then determine the resulting CAs with corresponding neurons, and retroactively sort the neurons into the CA-groups. Bottom row: average synaptic weight of recurrent synapses within the corresponding groups of neurons (CA1, CA2, and other neurons RR). Before learning (test 0), no specific synaptic structures are present. After the first learning phase (test 1), the first group of neurons becomes strongly interconnected, thus, a CA (CA1), which becomes also strongly connected to the active input population I1. (test 2) The second learning phase yields the formation of a second CA (CA2), which is linked to the second input population I2. (B): The formation of CAs is also indicated by the reduction of the average shortest path length (ASPL) between stimulus-activated neurons in the memory area. (Error bars are small and overlapped by symbols). (C): After both learning phases, the response vector overlap (RVO) between neurons activated by S1 and activated by S2 depends non-linearly on the disparity between the stimulus patterns. (A-C): Data presented are mean values over 100 repetitions. Explicit values of mean and standard deviation are given in Supplementary Table S1

**Figure 3.**
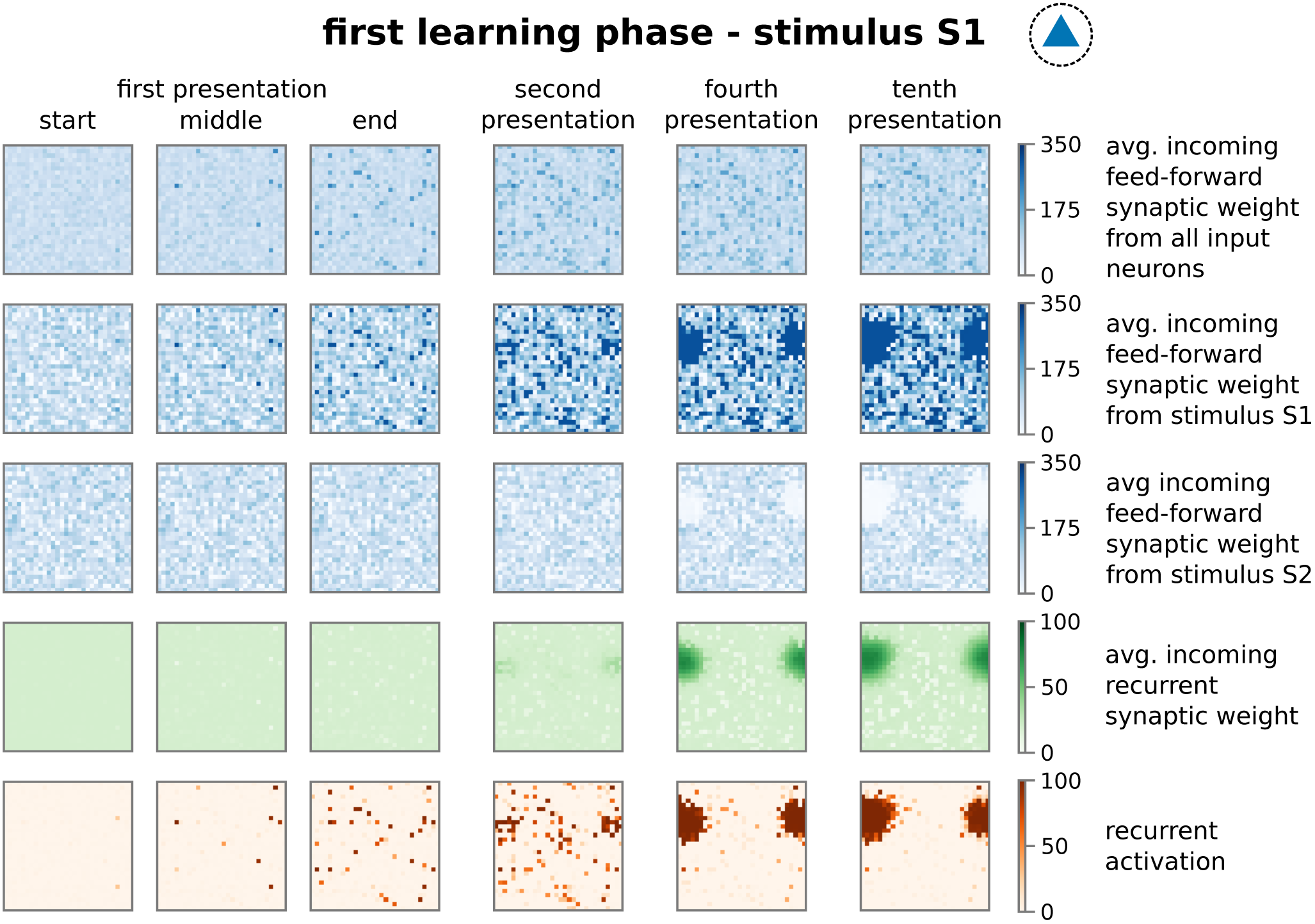
The repetitive presentation of a stimulus (here S1) triggers changes in feed-forward and recurrent synaptic weights as well as neural activities resulting to the proper formation and allocation of a CA. Each panel represents properties of the recurrent network in the memory area with neurons ordered on a 30*x*30 grid as indicated in Supplementary Figure S1A at different points of the protocol during the first learning phase. Thus, each dot in a panel represents a property of a neuron in the memory area, which is connected to the neighbouring dots as shown in Supplementary Figure S1 B. These properties are; first row: average feed-forward synaptic weights from all input neurons; second row: average feed-forward synaptic weights from the subset of I1-input neurons; third row: average feed-forward synaptic weights from the subset of I2-input neurons; forth row: average incoming synaptic weight from neurons of the memory area (recurrent synapses); fifth row: firing rate of the corresponding neuron. Please note that we consider torus-like periodic boundary conditions.

### Formation and allocation of a second memory representation

After showing that the synaptic dynamics of the feed-forward connection does not impede the formation of a CA as internal representation of a stimulus, next, we will demonstrate that the recurrent dynamics (thus, a formed CA) does not obstruct the feed-forward dynamics given new stimuli. Clearly the presence of a memory representation can alter the self-organizing dynamics shown before, which could impede the proper formation and allocation of representations of further stimuli. For instance, the existence of a CA in the neuronal network could bias the adaptations of the feed-forward synapses such that a new stimulus is also assigned to this CA. This would imply that the neural circuit is unable to discriminate between the originally CA-associated stimulus and the new stimulus. Thus, to investigate the influence of prior learning, we repeatedly present a second, different stimulus S2 (second learning phase; 1 C) after the proper formation of the CA associated to stimulus S1 and analyse whether a second CA is formed which is independent of the first one. Similar to the first learning phase, the repeated presentation of stimulus S2 (here, stimulus-associated activity patterns in the input area I1 and I2 have a *stimulus disparity* equals 1 indicating no overlap between patterns; see Supplementary Section A) yields via activity pattern I2 in the input area the activation of a group of interconnected neurons in the memory area (decreased ASPL; 2 B, test 2, red). In addition, the stimulation triggers a strengthening of the corresponding recurrent synaptic weights (2 A, bottom row, test 2, CA2; 4). Thus, the stimulus-dependent formation of a new CA is not impeded by the existence of another CA. Furthermore, both CAs are distinguishable as they do not share any neuron in the memory area (2 C, disparity equals 1; 4, forth row, tenth presentation; Supplementary Figure S3). As indicated by the response vector overlap (RVO; basically the number of neurons activated by both stimuli), this depends on the disparity between stimuli; for quite dissimilar stimuli both CAs are separated (disparity ≳ 0.5 yields RVO ≈ 0), for more similar stimuli the system undergoes a state transition (0.3 ≲ disparity ≲ 0.5 yields RVO > 0), and for quite similar stimuli both stimuli activate basically the same group of neurons (disparity ≲ 0.3 yields RVO > 100 given that 120 ± 4 neurons are on average part of a CA; Supplementary Figure S4). Note that the latter demonstrates that the network does correctly assign noisy versions of a learned stimulus pattern (*pattern completion*^15^) instead of forming a new CA, while the first case illustrates that the network performs *pattern separation*^15^ to distinguish different stimuli. This indicates a correct assignment of the stimuli to the corresponding CAs, such that also in the presence of another CA the weight changes of synapses between input pattern and newly formed CA are adapted accordingly (2 A, test 2, I2 to CA2). Thus, the self-organizing dynamics yields the formation and allocation of a new CA during the second learning phase. Note that the synaptic weights of the initially encoded CA are not altered significantly during this phase (2 A). But, although stimulus S1 is not present, the second learning phase leads to a weakening of synapses projecting from corresponding input neurons I1 to the newly formed CA considerably below control (2 A, test 2, I2 to CA1). Similarly, during the first learning phase the synaptic weights between input neurons I2 and the first cell assembly are also weakened (2 A, test 1, I2 to CA1). Apparently, this weakening of synapses from the other, non-assigned stimulus to a CA reduces the chance of spurious activations. In addition, as we will show in the following, this weakening is also an essential property enabling the proper functioning of the neural system. In summary, these results of our theoretical network model show that the self-organized processes resulting from the interplay between synaptic plasticity and scaling coordinates the synaptic and neuronal dynamics such that the proper formation and allocation of several memory representations without significant interferences between them is enabled.

**Figure 4.**
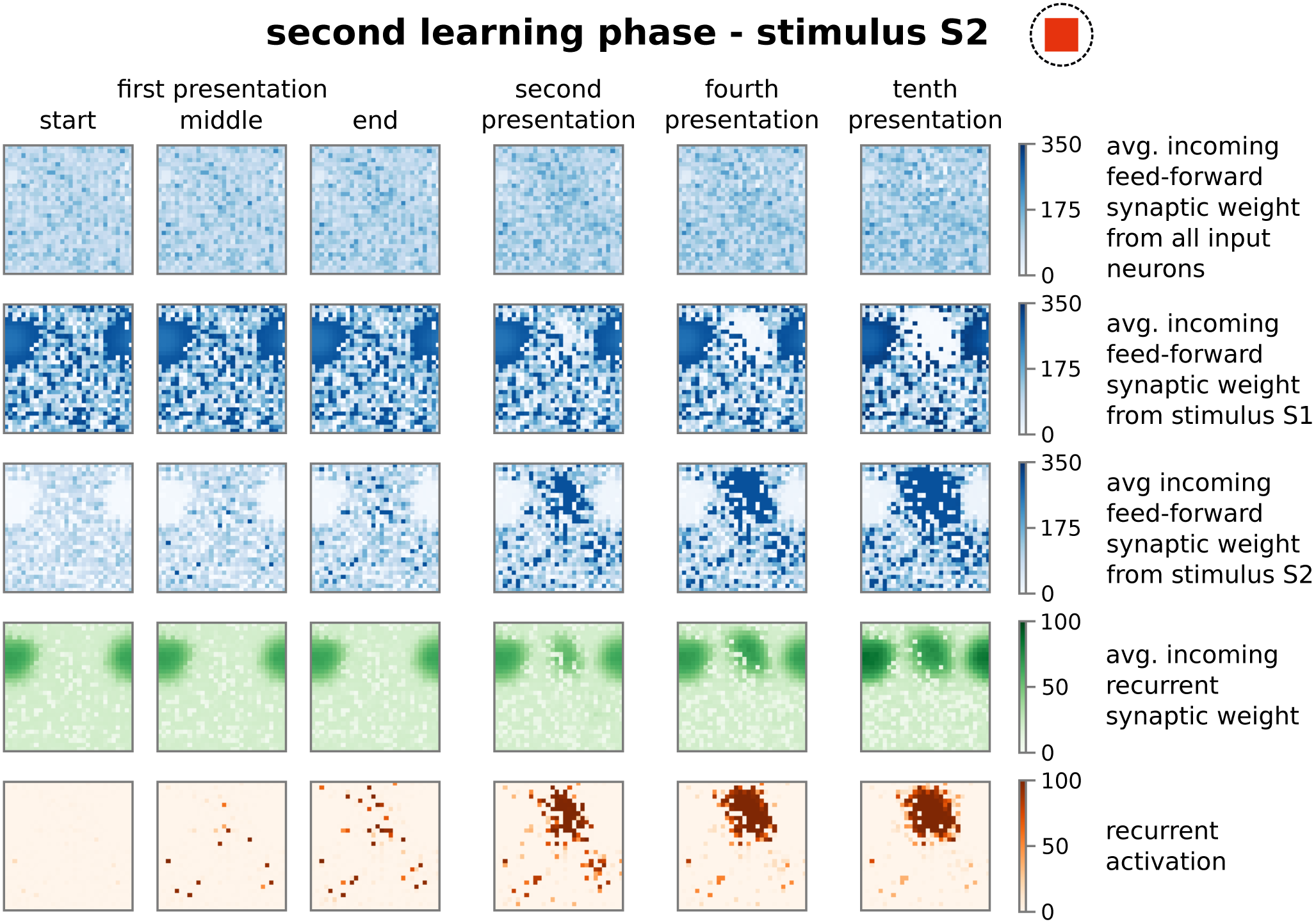
The presentation of a second stimulus (here S2) yields the formation of a second, distinct group of strongly interconnected neurons or CA. The structure of the sub-plots is the same as in 3. The first learning phase yields the encoding of a highly interconnected sub-population of neurons in the memory area (2 & 3). However, due to the interplay between Hebbian synaptic plasticity and synaptic scaling this CA cannot be activated by the second stimulus S2. Instead, the process of initially scattered activation (dark red dots in the fifth row; first presentation) and the following neuronal and synaptic processes (fourth to tenth presentation), as described before, are repeated yielding the formation of a second CA representing stimulus S2. Please note that both representations do not overlap (see Supplementary Figure S3).

### Generic properties of synaptic adaptations required for the formation and allocation of memory representations

In order to obtain a more detailed understanding of the self-organizing coordination of synaptic and neuronal dynamics by synaptic plasticity and scaling underlying the reliable formation and allocation of memory representations, we have to reduce the complexity of the model to enable (partially) analytical investigations. As already indicated by the above shown results (2 A), the main features of the self-organizing network dynamics can be described by considering the synaptic weights averaged over the given neuronal populations. Thus, we assume that the different involved neuronal populations in the input (input-pattern I1 and I2 neurons) and memory area (populations CA1 and CA2 becoming CAs in the memory area) are by themselves homogeneous allowing the derivation of a theoretical model describing the average population dynamics (5 A). For this, we combine the neuronal dynamics of all neurons within such a population (groups of I1-, I2-, CA1-, CA2-neurons) and describe them by the average neuronal dynamics of the population such that we obtain four variables each describing the average firing rate of one population or group of neurons (I1: *Ī*_1_; I2: *Ī*_2_; CA1: *F̄*_1_; CA2: *F̄*_2_). Similarly, we combine the synaptic dynamics of all synapses to describe them by the average synaptic dynamics for all connections within a neuronal population (within CA1: 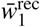; within CA2: 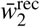) and for all connections between populations (from I1 to CA1: 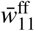; from I1 to CA2: 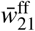; from I2 to CA1: 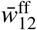; from I2 to CA2: 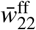). As activities and weights of the remaining neurons and synapses (IR and RR groups of neurons in 1 D) remain small, in the following, we neglect their influence on the system dynamics. The inhibition is considered similarly (see *Materials and Methods*) and the values of some system parameters are taken from full network simulations (Supplementary Figure S4). Please note, we re-scale the average neuronal activities and the average synaptic weights such that, if the weights equal one, learning is completed (see *Materials and Methods*). By considering the average neuronal and synaptic dynamics, we map the main features of the complex network dynamics with ≈ *N*2 dimensions (*N* is the number of neurons in the network) to a 8-dimensional theoretical model. Given such a population model of our adaptive network, first, we investigate the formation of a memory representation in a blank, random neural circuit (thus, *Ī*_1_ > 0 and *Ī*_2_ = 0). As mentioned before, the advantage of using a population model is the reduced dimensionality enabling, amongst others, the analytical calculation of the nullclines of the system to classify its dynamics. Nullclines show states of the system in which the change in one dimension equals zero^43, 44^. Thus, the intersection points of all nullclines of a system are the fixed points of the systems dynamics describing the system states in which no change occurs. Such fixed points can either be stable (attractive), thus the system will tend to converge into this point or state, or unstable (repulsive) meaning that the system will move away from this state (Supplementary Section C). Such dynamics can be summarized in a phase space diagram, in which each point indicates a possible state of the system and we can determine from the relative position of this state to the nullclines and fixed points to which new state the system will go to. The population model of the network model has 8 dimensions, thus, 8 nullclines and a 8-dimensional phase space (5 A); however, by inserting solutions of some nullclines into others (Supplementary Section B), we can project the dynamics onto two dimensions for visualization. In the following, we project the dynamics onto the average recurrent synaptic weights of both populations in the memory area (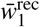, 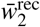) to investigate the dynamics underlying the formation of a CA during the first learning phase (5 B, top; please see 5 B, bottom, for corresponding activity levels).

**Figure 5.**
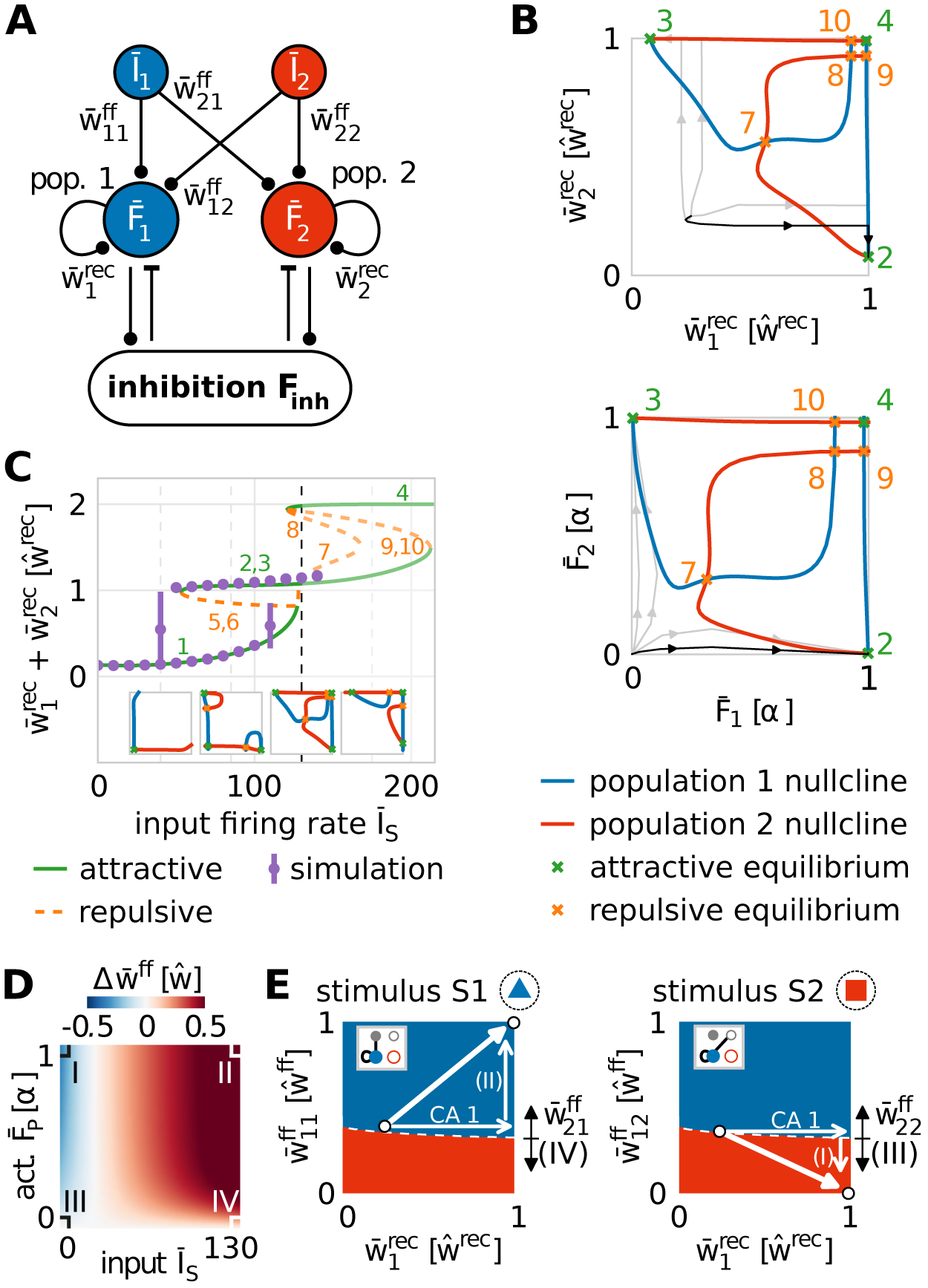
Population model of network dynamics enables analytical derivation of the underlying neuronal and synaptic dynamics. (A): Schema of the population model for averaged network dynamics (bars above variables indicate the average over all neurons in the population. *Ī_s_*: firing rate of input population *s* ∈ {1, 2}; *F̄_p_*: neural activity of population 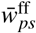: weight of feed-forward synapses from population 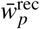: weight of recurrent synapses within population *p*. (B): The intersections of the population nullclines projected into weight (top) and activity (bottom) space reveal several fixed points (attractive: green; repulsive: orange) indicating the formation of a CA (green markers 2 and 3), if the system deviates from the identity line. Numbers correspond to labels of fixed points. (C): The bifurcation diagram (labels as in (B) and insets) of the network indicates that CAs are formed for a wide variety of input amplitudes (*Ī_A_* ≲ 120). The dashed line illustrates the value used in (B). Solutions of the full network model (purple dots) and population model match. (D): The dynamics of feed-forward synaptic weights depends on the firing rate of the input population and of the population in the memory area. There are four different cases (I-IV) determining the system dynamics. (E): These cases (indicated by arrows with Roman numbers: I-IV) together with the potentiation of recurrent synapses (arrow labelled CA1) yield the self-organized formation and allocation of CAs. Namely, during the first learning phase, synaptic changes drive the system (white dot) into regimes where either population 1 (blue) or population 2 (red) will represent the presented stimulus (left: stimulus S1; right: stimulus S2). Details see main text.

Thus, the projection of the solutions of the nullclines of the system dynamics onto the average recurrent synaptic weights of the two neuronal groups shows that the recurrent dynamics during the first learning phase are dominated by three fixed points: one is unstable (orange, 7; more specifically, it is a saddle point) and two are stable (green, 2 and 3). As the recurrent synaptic weights before learning are in general rather weak (which "sets" the initial state in this diagram), the fixed points 4, 8, 9, 10 cannot be reached by the system and, thus, they do not influence the here discussed dynamics. The two stable fixed points represent that one of the two neuronal populations becomes a strongly interconnected CA, while the other remains in a weakly interconnected state. For instance, in state 2 the first population is a CA as it is strongly interconnected 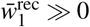 and the second population of neurons remains weakly connected (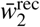 has only about 10% of the value of 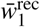). The unstable or repulsive fixed point lies on the identity line 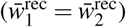 having the same distance to both stable, attractive states. The resulting mirror symmetry in the phase space implies that the dynamics on the one side of the identity line, reaching the stable fixed point lying on this side, equals the dynamics on the other side. Please note that the form of the nullclines and, thus, the existence and positions of the fixed points of the system dynamics depend on the mechanisms determining the synaptic adaptations. In other words, given a strong stimulus, the interplay between synaptic plasticity and scaling coordinates the recurrent synaptic dynamics such that the system by itself has to form a CA (reach one of both stable states). The question, which of both stable states is reached, translates into the question, to which group of neurons will the stimulus be allocated. As both groups are before learning quite similar, the initial state of the system will be close to the identity line. Only a minor difference between both groups (e.g., slightly different recurrent synaptic weights or a little different number of feed-forward synapses) results to a small variation in the initial condition of the first learning phase such that the system is slightly off the identity line (see traces for examples). Given the difference, the system will be on one side of the identity line and converge to the corresponding fixed point implying that the corresponding group of neurons will become the internal representation (e.g., black trace). Note that this symmetry-dependent formation of a CA is quite robust as long as the input firing rate is above a certain threshold (*Ī_A_* ≳ 120), which agrees with results from the more detailed network model discussed before (purple dots; 5 C). Below this threshold, the system remains in a state both groups of neurons are not becoming CAs (state 1 in 5 C; see also first two insets). Thus, the existence of the threshold predicts that a new, to-be-learned stimulus has to be able to evoke sufficient activity in the input area (above the threshold) to trigger the processes of memory formation; otherwise, the system will not learn.

In parallel to the development of the recurrent synaptic weights, the synaptic weights of the feed-forward connections change to assure proper memory allocation. Thus, we derive analytically the activity-dependency of the interaction of synaptic plasticity and scaling and obtain the change of the feed-forward synaptic weights (∆*w̄*^ff^), expected during the first learning phase, given different activity conditions of the input (*Ī_s_*) and CA-populations (*F̄_p_*, *s, p* ∈ {1, 2}; 5 D and Supplemenatry Section C). As expected, the combination of both activity levels for a certain duration determines whether the weights of the feed-forward synapses are potentiated (red), depressed (blue), or not significantly adapted (white). In general, if both activities are on a quite high level, synapses are potentiated (case II; so-called homosynaptic potentiation;^45^). If the pre-synaptic activity (input population) is on a low level and the post-synaptic activity (CA-population) is on a high level, on average, feed-forward synapses are depressed (case I; so-called heterosynaptic depression;^45^). However, if the post-synaptic activity is low, synaptic changes are negligible regardless of the level of pre-synaptic activity (cases III and IV).

The different parts of recurrent and feed-forward synaptic dynamics, described before, together lead to the formation and allocation of a CA as described in the following. For this, given the presentation of a to-be-learned stimulus, we have to consider the basins of attraction of the system in the phase space (5 E) projected onto different types of connections (insets; Gray indicates the active input population). If the system is in a certain state, we marked by the colour of this state which group of neurons will become a CA and will be assigned to the stimulus presented (left: S1 is presented; right: S2 is presented). The mirror symmetry described before (5 B) maps to the boundary (white dashed line in 5 E) between both basins of attraction (blue: population 1 becomes the internal representation; red: population 2 becomes the CA). Thus, during the first learning phase (stimulus S1; 5 E, left), a small variation in initial conditions breaks the symmetry such that the system is, in the example highlighted in 5 B (black trace), in the basin of attraction of population 1 becoming the internal representation (dot nearby symmetry line in blue area). This leads to the strengthening of the recurrent synapses within population 1 forming a CA (increase of 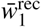; 5 B, E). In parallel, the synaptic strengthening induces an increase of the activity level of the population (*F̄*_1_ black trace in 5 B, bottom) yielding, together with the high activity level of input population I1 (Ī≫0), an average increase of the corresponding feed-forward synapses (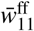; case II in 5 D). These synaptic changes push the system further away from the symmetry condition (white arrows; 5 E, left) implying a more stable memory representation. Note that changing the strength of synapses connecting input population I1 with population 2 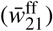 could result in a shift of the symmetry condition (indicated by black arrows) counteracting the stabilization process. However, this effect is circumvented by the system, as the second population has a low activity level and, therefore, corresponding feed-forward synapses are not adapted (case IV in 5 D). Thus, during the first learning phase, the formation and allocation of an internal representation is dictated by the subdivision of the system phase space into different basins of attraction of the stable fixed points such that small variations in the before-learning state of the network predetermines the dynamics during learning. This subdivision, in turn, emerges from the interplay of synaptic plasticity and scaling.

How do these synaptic and neuronal dynamics of the allocation and formation of the first CA influence the dynamics of the second learning phase? In general, the formation of a CA acts as a variation or perturbation of the initial condition breaking the symmetry for the second learning phase (stimulus S2; 5 E, right). The formation of the first CA pushes the system into the blue area (CA1-arrow). This indicates that, if stimulus S2 is presented, the feed-forward synapses would be adapted such that population 1 would also represent stimulus S2. This would impede the discrimination ability of the network between stimulus S1 and S2. However, during the first learning phase, as the input population I2 of stimulus S2 is inactive, synapses projecting from I2-input neurons to population 1 neurons are depressed (case I in 5 D; downward arrow in 5 E, right) and the system switches into the red area. This area indicates that, if stimulus S2 is presented as during the second learning phase, population 2 would form a CA representing stimulus S2 and not population 1. Please note that this switch can be impeded by adapting the connections from the I2-input population to population 2 during the first learning phase 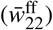 shifting the symmetry condition (black arrows in 5 E, right). But, similar to before, this effect is circumvented by the system, as population 2 is basically inactive resulting to case III (5 D). Thus, after the first learning phase, the synaptic dynamics regulated by the combination of Hebbian plasticity and synaptic scaling drives the system into an intermediate state, which implies that the system will definitely form a new CA during the second learning phase (see 6 for further phase space projections and dynamics during second learning phase). These results indicate that this intermediate state can only be reached if synaptic adaptations comprise three properties implied by the four cases I-IV (5 D): (i) homosynaptic potentiation (case I), (ii) heterosynaptic depression (case II), and (iii) the down-regulation of synaptic weight changes by the post-synaptic activity level (cases III and IV).

**Figure 6.**
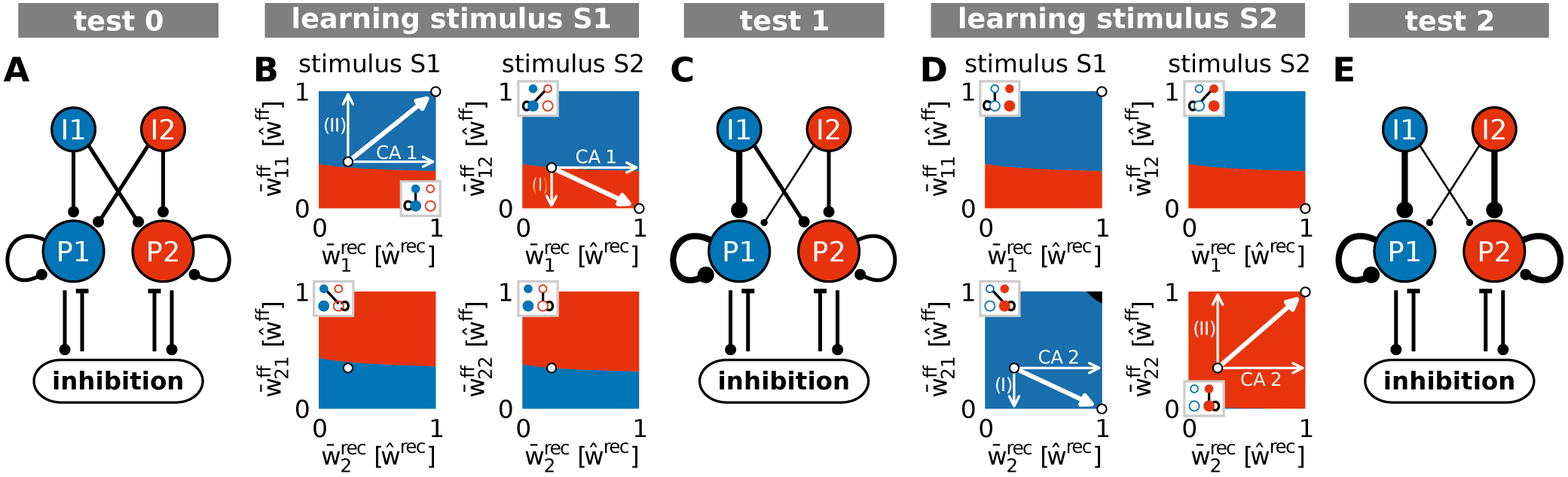
Summary of the synaptic changes and their implication on the formation and allocation of memory representations. The interaction of synaptic plasticity and scaling brings a blank network (A; test 0) during the first learning phase (B) to an intermediate state (C; test 1). From this intermediate state, the second learning phase (D) yields the system into the desired end state (E; test 2), in which each stimulus is allocated to one CA (*S*1/*I*1 to pop. 1/CA1 and *S*2/*I*2 to pop. 2/CA2). (A), (C), (E): Thickness of lines is proportional to average synaptic weight. (B), (D): Similar to panels in 5 E. Black area indicates regimes in which both populations would be assigned to the corresponding stimulus.

### Modelling experimental findings of competition-based memory allocation

In addition to the described three properties, the here-proposed model implies the existence of a symmetry condition underlying the formation and allocation of memory representations. Small variations in the initial condition of the system suffice to break this symmetry. These variations could be, aside from noise, enforced experimentally by adapting neuronal parameters in a local group of neurons. Amongst others, several experiments^42, 46^ indicate that the probability of a group of neurons to become part of a newly formed memory representation can be influenced by changing their excitability pharmacologically (e.g., by varying the CREB concentration). We reproduced such manipulations in the model by adapting the neuronal excitability of a group of neurons accordingly and analysing the data similar to experiments (see *Materials and Methods*). Thus, we investigated the probability of a single neuron to become part of a CA averaged over the whole manipulated group of neurons (relative recruitment factor) and compared the results to experimental findings (7 A).

On the one hand, if the excitability of a group of neurons is artificially increased briefly before learning, the probability of these neurons to become part of the memory representation is significantly enhanced. On the other hand, if the excitability is decreased, the neurons are less likely to become part of the representation. Considering the theoretical results shown before (5 B), this phenomenon can be explained as follows: the manipulation of the excitability in one population of neurons changes the distance between the repulsive state (orange; 7 B) to the two attractive states (green). Thus, an increased (decreased) excitability yields a larger (smaller) distance between the repulsive state and the attractive state related to the manipulated population (e.g., instance iii for increased excitability of population 1). This larger (smaller) distance implies a changed basin of attraction of the manipulated population enhancing the chance that the initial condition of the network (black dots) lies within this basin. This implies an increase (decrease) of the probability that this group of neurons becomes a CA, as depicted by the variation of the experimentally measured single neuron probability. In addition, the theoretical analysis yields the prediction that the measured effects will be altered by manipulating other parameters. For instance, if the synaptic weight of the population with increased excitability is on average decreased before stimulus presentation (e.g. by PORCN;^47^), the network’s initial condition is shifted such that the CREB-induced influence on the relative recruitment factor is counterbalanced (7 A, instance iv). Thus, the match between model and *in-vivo* experimental data supports the here-proposed hypothesis that the combination of Hebbian synaptic plasticity and synaptic scaling coordinates the synaptic and neuronal dynamics enabling the self-organized allocation and formation of memory representations.

**Figure 7.**
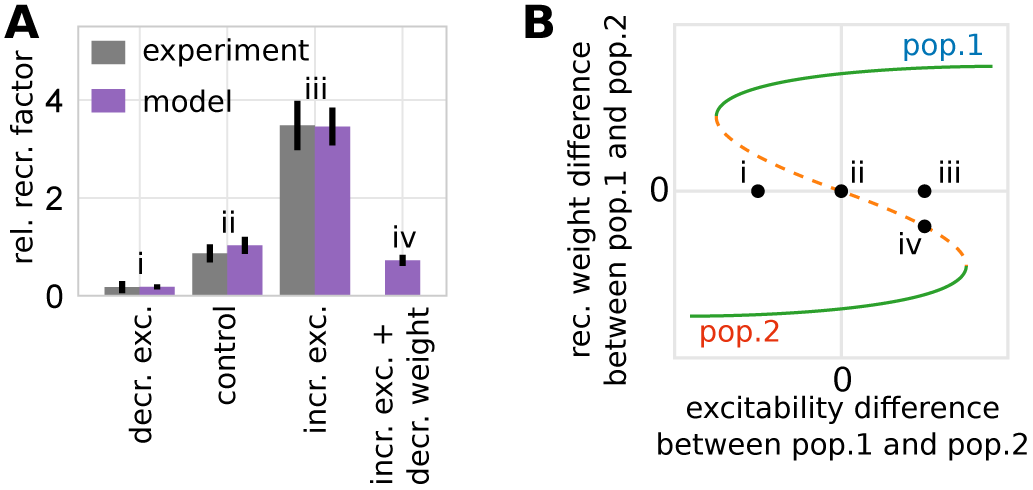
The model of synaptic plasticity and scaling matches experimental *in-vivo* data and provides experimentally verifiable predictions. (A): The artificial modification of the excitability of a subset of neurons alters the probability of these neurons to become part of a memory representation (normalized to control). Experimental data are taken from^42^. Data presented are mean values with standard error of the mean. Labels correspond to instances shown in (B). (B): The alteration of the excitability of one group of neurons (here pop. 1) compared to others yields a shift of the positions of the system’s fixed points and basins of attractions (here shown schematically; for details see Supplementary Figure S6) inducing a bias towards one population (e.g., from instance ii to iii by increasing the excitability of population 1). (A), (B): The model analysis yields the prediction that this excitability-induced bias can be counterbalanced by, for instance, additionally decreasing the average synaptic weight of the manipulated population before learning (here by a factor of 0.1). This additional manipulation shifts the initial state of the network back to the symmetry condition (orange; instance iv).

## Discussion

Our theoretical study indicates that the formation as well as the allocation of memory representations in neuronal networks depend on the self-organized coordination of synaptic changes at feed-forward and recurrent synapses. We predict that the combined dynamics of conventional Hebbian synaptic plasticity and synaptic scaling could be sufficient for yielding this self-organized coordination as it implies three generic properties: (i) homosynaptic potentiation, (ii) heterosynaptic depression, and (iii) the down-regulation of synaptic weight changes by the post-synaptic activity level.

Previous theoretical studies show that the properties (i) and (ii) are required in recurrent neuronal networks to dynamically form memory representations^23–25^. However, these studies do not consider the feed-forward synaptic dynamics. On the other hand, studies analysing feed-forward dynamics, such as the self-organization of cortical maps^34–36^, also indicate the importance of homosynaptic potentiation and heterosynaptic depression. However, these studies do not consider the recurrent synaptic dynamics. Only by considering both feed-forward *and* recurrent synaptic dynamics, we revealed the requirement of property (iii) that a low level of post-synaptic activity curtails the synaptic changes which is also supported by experimental evidence^48, 49^.

Note that property (iii) is realized by both mechanisms: Hebbian synaptic plasticity as well as synaptic scaling. By contrast, property (i) is implemented by Hebbian synaptic plasticity only and property (ii) is realized by synaptic scaling only. This indicates that synaptic scaling could have an essential role in the allocation and formation of multiple memory representations beyond the widely assumed stabilization of neural network dynamics^7, 38, 40, 50, 51^.

Similar to previous studies^23, 26, 41^, we consider here an abstract model to describe the neuronal and synaptic dynamics of the network. Despite the abstract level of description, the model matches experimental *in-vivo* data of memory allocation. Other theoretical models match similar experimental data^32, 52^; however, these models are of greater biological detail including more dynamic processes (e.g., short-term plasticity). However, only by considering an abstract model, we have been able to derive analytical expressions such that we could find the requirement of the three generic properties yielding the proper formation and allocation of memories. Remarkably, the synaptic plasticity processes considered in the detailed models^32, 52, 53^ also imply the three generic properties (i-iii) supporting our findings. Further investigations are required to assess possible differences between different realizations of the three generic properties. For this, the here used theoretical methods from the field of non-linear dynamics^43, 44^ seem to be promising given their ability to derive and classify fundamental system dynamics, which can be verified by experiments.

Our results indicate that the combined dynamics of Hebbian synaptic plasticity and synaptic scaling are sufficient to coordinate the synaptic and neuronal dynamics underlying the allocation and formation of memory representations. Interestingly, for several different stimuli the dynamics always yields the formation of separated memory representations. In particular the process of heterosynaptic depression impedes the formation of overlaps between several memory representations (neurons encoding several stimuli). However, amongst others, experimental results indicate that memory representations can overlap^11, 54, 55^ and, in addition, theoretical studies show that overlaps increase the storage capacity of a neuronal network^19^ and can support memory recall^56^. To partially counterbalance the effect of heterosynaptic depression to enable the formation of overlaps, further time-dependent processes are required. For instance, the CREB-induced enhancement of neuronal excitability biases the neuronal and synaptic dynamics such that the respective subgroup of neurons is more likely to be involved in the formation of a memory representation (7;^42, 52^). Furthermore, the dynamics of CREB seem to be time-dependent^32, 42, 46^. Therefore, the enhancement of CREB can counterbalance heterosynaptic depression for a given period of time and, by this, could enable the formation of overlaps. We expect that the impact of such time-dependent processes on the dynamics of memories can be integrated into the here-proposed model to analyse the detailed formation of overlaps between memory representations. Thus, our study shows that the interplay between synaptic plasticity and scaling is required to include all three generic properties of synaptic adaptation enabling a proper formation and allocation of memories. In addition, given the here-derived theoretical model, other mechanisms can be included to investigate systematically their functional implication on the self-organized, complex system dynamics underlying the multitude of memory processes.

## Methods

### Numerical simulations

The neuronal system in the numerical model consist of 936 excitatory neurons (36 in input area, 900 in memory area) and a single inhibitory unit. The inhibitory unit describes a population of inhibitory neurons which are connected to the excitatory neurons in an all-to-all manner. Given the long time scales considered in this study, all neurons are described by a rate-coded leaky integrator model. The memory area is arranged as a quadratic neural grid of 30×30 units. Each neuron within the grid receives excitatory connections from four randomly chosen input neurons. In addition, it is recurrently connected to its circular neighbourhood of radius four (measured in neuronal units; for visualization see 1 B and Supplementary Figure S1) and to the global inhibitory unit. Initially, recurrent synaptic weights of existing connections equal to 0.25 • *ŵ*^rec^ and feed-forward synaptic weights are drawn from a uniform distribution {0, 0.7 *• ŵ*^ff^} 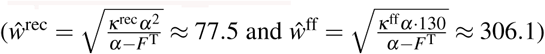. Connections to and from the inhibitory neuron are at fixed and homogeneous weight (for detailed values see 1).

#### Neuron model

The membrane potential 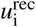 of each excitatory neuron *i* in the memory area is described as follows:

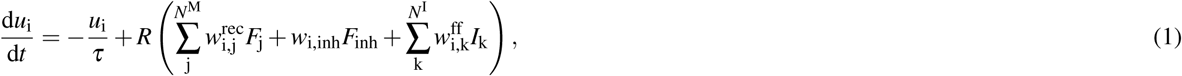

with the membrane time constant *τ*, membrane resistance *R*, number of neurons in the memory area *N*^M^, number of neurons in the input area *N*^I^, and firing rate *I*_k_ of input neuron *k*. The membrane potential is converted into a firing rate *F*_i_ by a sigmoidal transfer function with maximal firing rate *α*, steepness *β* and inflexion point *ε*:

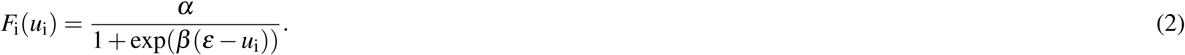

The global inhibitory unit is also modelled as a rate-coded leaky integrator receiving inputs from all neurons of the memory area. Its membrane potential *u*_inh_ follows the differential equation 3 with inhibitory membrane time scale *τ*_inh_ and resistance *R*_inh_. The potential is converted into a firing rate *F*_inh_ by a sigmoidal transfer function (equation 4):

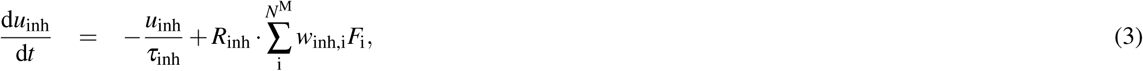

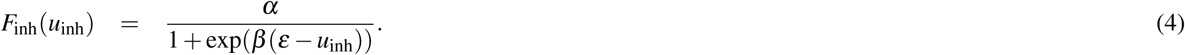

As the neurons in the input area form the stimuli, their output activation is set manually. Thus no further description is needed.

#### Synaptic plasticity

The weight changes of the excitatory feed-forward (equation 5) and recurrent synapses (equation 6) are determined by the combined learning rule of conventional Hebbian synaptic plasticity (first term) and synaptic scaling (second term) with time constants *μ*, *κ*^ff^, *κ*^rec^, and target firing rate *F*^T^. The differential equation for the synaptic weight of a feed-forward connection 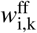 from input neuron *k* ∈ {0, ⋯, *N*^I^} to memory neuron *i* ∈ {0, ⋯, *N*^M^} is:

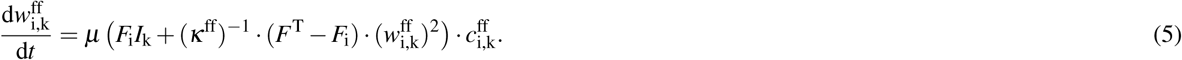

The dynamics of the synaptic weight of a recurrent connection 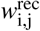 from memory neuron *j* ∈ {0, ⋯ *N*^M^} to memory neuron *i* ∈ {0, ⋯ *N*^M^} is determined by:

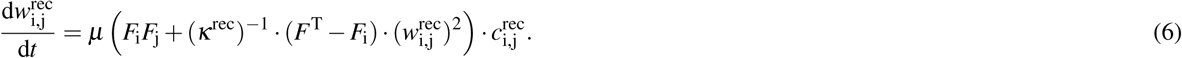

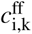 and 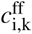 are the entries in the feed-forward and recurrent connectivity matrices of value 1, if the connection exists, or otherwise of value 0.

All other connections are considered to be non-plastic.

**Table 1.** Model Parameters: Variable, Descriptions and used Values

| Variable | Description | Value |
| --- | --- | --- |
| $\tau$ | membrane time constant (memory area) | 0.01 |
| $R$ | membrane resistance (memory area) | 1/11 |
| $N^M$ | number of neurons in memory area | 900 |
| $N^I$ | number of neurons in input area | 36 |
| $I_k$ | input rate | {0,130} |
| $\alpha$ | maximum firing rate | 100 |
| $\beta$ | sigmoid steepness | 0.05 |
| $\varepsilon$ | sigmoid inflexion point | 130 |
| $\mu$ | plasticity time constant | 1/15 |
| $F^T$ | target firing rate | 0.1 |
| $\kappa^{\text{rec}}$ | scaling time constant (recurrent) | 60 |
| $\kappa^{\text{ff}}$ | scaling time constant (feed-forward) | 720 |
| $\tau_{\text{inh}}$ | membrane time constant (inhibitory unit) | 0.02 |
| $R_{\text{inh}}$ | membrane resistance (inhibitory unit) | 1 |
| $w_{\text{inh},i}$ | synaptic weight to inhibitory unit | 0.6 |
| $w_{i,\text{inh}}$ | synaptic weight from inhibitory unit | 1200 |

#### Coding framework

The differential equations have been solved numerically with the Euler method with a time step of 5 ms using Python 3.5.

#### Simulation of self-organized formation of two CAs

The system undergoes two learning phases presenting two completely dissimilar input patterns I1 (first phase) and I2 (second phase). For input pattern I1 half of the input neurons are set to be active at 130 Hz whereas the other half remains inactive at 0 Hz and vice versa for input pattern I2. During a learning phase the respective pattern is presented 10 times for 5 sec with a 1 sec pause in between. Both learning phases are embraced by test phases in which plasticity is shut off and both patterns are presented for 0.5 sec each to apply measures on the existing memory structures.

#### Comparison to experimental data

The manipulation of the neuronal excitability has been done by adapting the value for *ε* in the transfer function of the neuron model, i.e. shifting its inflexion point to lower (increased excitability) or higher (decreased excitability) values. Similar to the methods used in experiments^42^, we manipulated a sub-population of neurons within a randomly chosen circular area in the memory area (about 10% of the network). The relative recruitment factor is the relation of recruitment probabilities for manipulated and control neurons averaged over 100 repetitions.

### Population model

We consider two non-overlapping populations of *N* excitatory neurons and one inhibitory unit.

The state of every population *i* ∈ {1, 2} is determined by its mean membrane potential *ū_i_*, its mean recurrent synaptic weight 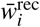 between neurons of the population, and the mean weight 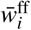 of feed-forward synapses projecting signals from the currently active input onto the population. We assume that the two populations interact solely through the inhibitory unit whose state is given by its membrane potential *u*_inh_. Thus, the dynamics of the model is described by a set of seven differential equations (see following section). To obtain its equilibria, we analytically derive the nullclines 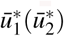 and 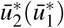 and numerically determine their intersections. The stability of an equilibrium is obtained from the sign of the eigenvalues of the system’s Jacobi Matrix.

For analyzing in which regimes population 1 and population 2 are assigned to an input stimulus as a function of the initial synaptic weights (5 E and 6), we initialize the system with the given combination of feed-forward and recurrent average synaptic weights and *ū*_1_ = *ū*_2_ = *ū*_inh_ = 0, simulate it for 100 sec, and assess which of the two populations is active. For further details and parameter values see *Supplementary Information*.

#### Model Definition

The two excitatory populations in the population model are described by their mean membrane potentials *ū_i_*, *i* ∈ {1, 2}, *k* ∈ {A, B}:

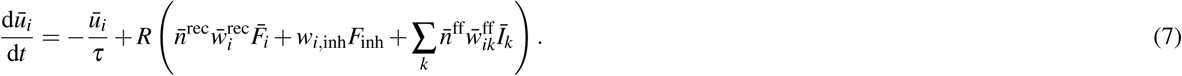

Here, the time scale *τ*, the resistance *R* and the synaptic weight *w_i,_*_inh_ have the same value as in the network simulations. The average number 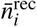 of incoming recurrent connections from neurons within the population as well as the number of feed-forward synapses transmitting signals from active inputs to every neuron (*n̄*^ff^) are taken from simulations (*n̄*^rec^ = 35, Supplementary Figure S4 C; *n*^ff^ = 2.3, Supplementary Figure S4 B).

The membrane potential of the inhibitory population is given by

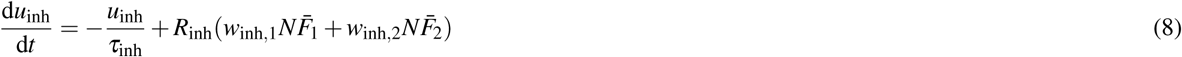

with *τ*_inh_, *R*_inh_ and *w*_inh,1_ = *w*_inh,2_ corresponding to the respective values in the network simulations. The number *N* of neurons per population is adjusted to the CAs in the network simulation and chosen as *N* = 120 (Supplementary Figure S3 A).

The transfer function of the neurons within the population is the same as for individual neurons:

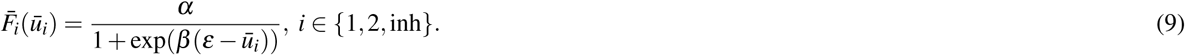

The synaptic weight changes of recurrent and feed-forward synapses follow the interplay of conventional Hebbian synaptic plasticity and synaptic scaling (*i* ∈ {1, 2}):

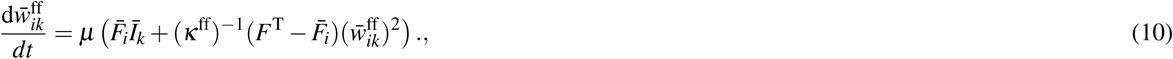

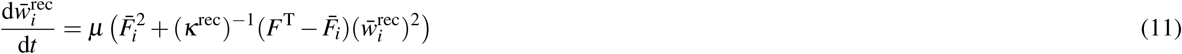

#### Data Display

The population model applies the same combined learning rule as the numerical simulation. We thus consider the memorization process completed when the dynamic fixed point of synaptic plasticity is reached, i.e. Hebbian plasticity and scaling compensate each other.

In order to focus the reader’s attention onto the populations dynamics as well as for illustrative reasons, we scale the explicit values of synaptic weights and activation in their depiction (5 B-E, 6 B,D, Supplementary Figure S6) with the respective values of their dynamic fixed points (weights; *ŵ*^ff^, *ŵ*^rec^) or the maximum value (activation; *α*). Hence, their values simply range from 0 to 1.

### Recruitment Basins

For determining the recruitment basins (5 E and 6), we exploit the symmetry of the system and that, in general, only one of the two stimuli (S1 or S2) is active. Accordingly, we approximate the second, inactive input to zero and neglect the respective feed-forward synapses. The population model is integrated with the given initial values of the feed-forward and recurrent weights and *ū*_1_ = *ū*_2_ = *ū*_inh_ = 0 for 100 s. At *t* = 100 s, we evaluate which of the two populations is active. 2 provides the exact used initial values.

**Table 2.** Initial values for Recruitment Basin Plots

| Figure | $\bar{u}_1$ | $\bar{u}_2$ | $\bar{u}_{inh}$ | $\bar{w}_{1A}^{ff}$ | $\bar{w}_{2A}^{ff}$ | $\bar{w}_1^{rec}$ | $\bar{w}_2^{rec}$ |
| --- | --- | --- | --- | --- | --- | --- | --- |
| 2E left; 2E right<sup>†</sup>;<br>3B top left, 3B top right<sup>†</sup>,<br>3B bottom right*<sup>†</sup>,<br>3D top left, 3D top right<sup>†</sup> | 0 | 0 | 0 | $0, \Delta w_{1A}^{ff}, \dots, \hat{w}_{1A}^{ff}$ | $0.35 \hat{w}_{2A}^{ff}$ | $0, \Delta w_1^{rec}, \dots, \hat{w}_1^{rec}$ | $0.25 \hat{w}_2^{rec}$ |
| 3B bottom left* | 0 | 0 | 0 | $0, \Delta w_{1A}^{ff}, \dots, \hat{w}_{1A}^{ff}$ | $0.40 \hat{w}_{2A}^{ff}$ | $0, \Delta w_1^{rec}, \dots, \hat{w}_1^{rec}$ | $0.25 \hat{w}_2^{rec}$ |
| 3D bottom left* | 0 | 0 | 0 | $0, \Delta w_{1A}^{ff}, \dots, \hat{w}_{1A}^{ff}$ | $1.00 \hat{w}_{2A}^{ff}$ | $0, \Delta w_1^{rec}, \dots, \hat{w}_1^{rec}$ | $1.00 \hat{w}_2^{rec}$ |
| 3D bottom right*<sup>†</sup> | 0 | 0 | 0 | $0, \Delta w_{1A}^{ff}, \dots, \hat{w}_{1A}^{ff}$ | $0.00 \hat{w}_{2A}^{ff}$ | $0, \Delta w_1^{rec}, \dots, \hat{w}_1^{rec}$ | $1.00 \hat{w}_2^{rec}$ |

\*) Using symmetry by commutating populations. †) Using symmetry by commutating inputs.

## Acknowledgements (not compulsory)

We thank Florentin Wörgötter for fruitful comments. The research was funded by the H2020-FETPROACT project Plan4Act (#732266) [JMA, CT], by the Federal Ministry of Education and Research Germany (#01GQ1005A, #01GQ1005B) [TN, CT], and by the International Max Planck Research School for Physics of Biological and Complex Systems by stipends of the country of Lower Saxony with funds from the initiative “Niedersächsisches Vorab” and of the University of Göttingen [TN].

## Author contributions statement

J.M.A. contributed the network simulations, T.N. contributed the population model. All authors reviewed the manuscript.

## Additional information

### Competing Interests

The authors declare no competing interests

## References

1. Martin, S. J., Grimwood, P. D. & Morris, R. G. M. Synaptic plasticity and memory: an evaluation of the hypothesis. Ann Rev Neurosci 23, 649–711, DOI: https://doi.org/10.1146/annurev.neuro.23.1.649 (2000)

2. Barbieri, F. & Brunel, N. Can attractor network models account for the statistics of firing during persistent activity in prefrontal cortex? Front Neurosci 2(1), 114–122, DOI: https://doi.org/10.3389/neuro.01.003.2008 (2008)

3. Palm, G., Knoblauch, A., Hauser, F. & Schüz, A. Cell assemblies in the cerebral cortex. Biol Cybern 108, 559–572, DOI: https://doi.org/10.1007/s00422-014-0596-4 (2014)

4. Takeuchi, T., Duszkiewicz, A. & Morris, R. The synaptic plasticity and memory hypothesis: encoding, storage and persistence. Philos Trans R Soc Lond B Biol Sci 369, 20130288, DOI: 10.1098/rstb.2013.0288 (2014)

5. Turrigiano, G. G., Leslie, K. R., Desai, N. S., Rutherford, L. C. & Nelson, S. B. Activity-dependent scaling of quantal amplitude in neocortical neurons. Nature 391, 892–896, DOI: https://doi.org/10.1038/36103 (1998)

6. Hengen, K. B., Lambo, M. E., Van Hooser, S. D., Katz, D. B. & Turrigiano, G. G. Firing rate homeostasis in visual cortex of freely behaving rodents. Neuron 80, 335–342, DOI: https://doi.org/10.1016/j.neuron.2013.08.038 (2013)

7. Tetzlaff, C., Kolodziejski, C., Timme, M. & Wörgötter, F. Synaptic scaling in combination with many generic plasticity mechanisms stabilizes circuit connectivity. Front Comput. Neurosci 5, 47, DOI: https://doi.org/10.3389/fncom.2011.00047 (2011)

8. James, W. The principles of psychology (New York: Henry Holt and Company, 1890)

9. Konorski, J. Conditioned reflexes and neuron organization (Cambridge: Cambridge University Press, 1948)

10. Hebb, D. O. The Organization of Behaviour (Wiley, New York, 1949)

11. Holtmaat, A. & Caroni, P. Functional and structural underpinnings of neuronal assembly formation in learning. Nat Neurosci 19, 1553–1562, DOI: https://doi.org/10.1038/nn.4418 (2016)

12. Bliss, T. V. P. & Lomo, T. Long-lasting potentiation of synaptic transmission in the dentate area of the anaesthetized rabbit following stimulation of the perforant path. J Physiol 232, 331–356, DOI: https://doi.org/10.1113/jphysiol.1973.sp010273 (1973)

13. Levy, W. B. & Steward, O. Temporal contiguity requirements for long-term associative potentiation/depression in the hippocampus. Neuroscience 8(4), 791–797, DOI: https://doi.org/10.1016/0306-4522(83)90010-6 (1983)

14. Malenka, R. C. & Bear, M. F. LTP and LTD: an embarrassment of riches. Neuron 44, 5–21, DOI: https://doi.org/10.1016/j.neuron.2004.09.012 (2004)

15. Hunsaker, M. R. & Kesner, R. P. The operation of pattern separation and pattern completion processes associated with different attributes or domains of memory. Neurosci Biobehav. Rev 37, 36–58, DOI: https://doi.org/10.1016/j.neubiorev.2012.09.014 (2013)

16. Hopfield, J. J. Neural networks and physical systems with emergent collective computational abilities. Proc Natl Acad Sci 79, 2554–2558, DOI: https://doi.org/10.1073/pnas.79.8.2554 (1982)

17. Hopfield, J. J. Neurons with graded response have collective computational properties like those of two-state neurons. Proc Natl Acad Sci 81, 3088–3092, DOI: https://doi.org/10.1073/pnas.81.10.3088 (1984)

18. Amit, D. J., Gutfreund, H. & Sompolinsky, H. Storing infinite numbers of patterns in a spin-glas model of neural networks. Phys. Rev. Lett. 55(14), 1530–1533 (1985)

19. Tsodyks, M. & Feigelman, M. Enhanced storage capacity in neural networks with low level of activity. Eur. Lett 6, 101–105 (1988)

20. Amit, D. J., Brunel, N. & Tsodyks, M. Correlations of cortical Hebbian reverberations: theory versus experiment. J Neurosci 14, 6435–6445, DOI: https://doi.org/10.1523/JNEUROSCI.14-11-06435.1994 (1994)

21. Buzsaki, G. Neural syntax, cell assemblies, synapsembles, and readers. Neuron 68, 362–385, DOI: https://doi.org/10.1016/j.neuron.2010.09.023 (2010)

22. Brunel, N. Is cortical connectivity optimized for storing information? Nat Neurosci 19, 749–755, DOI: https://doi.org/10.1038/nn.4286 (2016)

23. Tetzlaff, C., Kolodziejski, C., Timme, M., Tsodyks, M. & Wörgötter, F. Synaptic scaling enables dynamically distinct short- and long-term memory formation. PLoS Comput. Biol 9(10), e1003307, DOI: https://doi.org/10.1371/journal.pcbi.1003307 (2013)

24. Litwin-Kumar, A. & Doiron, B. Formation and maintaince of neuronal assemblies through synaptic plasticity. Nat Commun 5, 5319, DOI: https://doi.org/10.1038/ncomms6319 (2014)

25. Zenke, F., Agnes, E. J. & Gerstner, W. Diverse synaptic plasticity mechanisms orchestrated to form and retrieve memories in spiking neural networks. Nat Commun 6, 6922, DOI: DOI:10.1038/ncomms7922 (2015)

26. Tetzlaff, C., Dasgupta, S., Kulvicius, T. & Wörgötter, F. The use of Hebbian cell assemblies for nonlinear computation. Sci Rep 5, 12866, DOI: https://doi.org/10.1038/srep12866 (2015)

27. Rogerson, T. et al. Synaptic tagging during memory allocation. Nat Rev Neurosci 15, 157–169, DOI: https://doi.org/10.1038/nrn3667 (2014)

28. Willshaw, D. J., Buneman, O. P. & Longuet-Higgins, H. C. Non-holographic associative memory. Nature 222, 960–962, DOI:http://dx.doi.org/10.1038/222960a0 (1969)

29. Adelsberger-Mangan, D. M. & Levy, W. B. Information maintenance and statistical dependence reduction in simple neural networks. Biol Cybern 67, 469–477, DOI: https://doi.org/10.1007/BF00200991 (1992)

30. Knoblauch, A., Palm, G. & Sommer, F. T. Memory capacities for synaptic and structural plasticity. Neural Comput. 22(2), 289–341, DOI: https://doi.org/10.1162/neco.2009.08-07-588 (2010)

31. Babadi, B. & Sompolinsky, H. Sparseness and expansion in sensory representations. Neuron 83, 1213–1226, DOI: https://doi.org/10.1016/j.neuron.2014.07.035 (2014)

32. Kastellakis, G., Silva, A. J. & Poirazi, P. Linking memories across time via neuronal and dendritic overlaps in model neurons with active dendrites. Cell Reports 17, 1491–1504, DOI: https://doi.org/10.1016/j.celrep.2016.10.015 (2016)

33. Choi, J.-H. et al. Interregional synaptic maps among engram cells underlie memory formation. Science 360, 430–435, DOI: http://dx.doi.org/10.1126/science.aas9204(2018)

34. Sullivan, T. J. & de Sa, V. R. Homeostatic synaptic scaling in self-organizing maps. Neural Networks 19, 734–743, DOI: 10.1016/j.neunet.2006.05.006 (2006)

35. Stevens, J.-L. R., Law, J. S., Antolik, J. & Bednar, J. A. Mechanisms for stable, robust, and adaptive development of orientation maps in the primary visual cortex. J Neurosci 33(40), 15747–15766, DOI: https://doi.org/10.1523/JNEUROSCI.1037-13.2013 (2013)

36. Kohonen, T. Self-organized formation of topologically correct feature maps. Biol Cybern 43, 59–69, DOI: https://doi.org/10.1007/BF00337288 (1982)

37. Obermayer, K., Ritter, H. & Schulten, K. A principle for the formation of the spatial structure of cortical feature maps. Proc Natl Acad Sci 87, 8345–8349, DOI: https://doi.org/10.1073/pnas.87.21.8345 (1990)

38. Abbott, L. F. & Nelson, S. B. Synaptic plasticity: taming the beast. Nat Neurosci 3, 1178–1183, DOI: https://doi.org/10.1038/81453 (2000)

39. Gerstner, W. & Kistler, W. M. Mathematical formulations of Hebbian learning. Biol Cybern 87, 404–415, DOI: https://doi.org/10.1007/s00422-002-0353-y (2002)

40. Turrigiano, G. G. & Nelson, S. B. Homeostatic plasticity in the developing nervous system. Nat Rev Neurosci 5, 97–107, DOI: https://doi.org/10.1038/nrn1327 (2004)

41. Nachstedt, T. & Tetzlaff, C. Working memory requires a combination of transient and attractor-dominated dynamics to process unreliably timed inputs. Sci Rep 7, 2473, DOI: https://doi.org/10.1038/s41598-017-02471-z (2017)

42. Yiu, A. P. et al. Neurons are recruited to a memory trace based on relative neuronal excitability immediately before training. Neuron 83, 722–735, DOI: 10.1016/j.neuron.2014.07.017 (2014)

43. Glendinning, P. Stability, instability and chaos: An introduction to the theory of nonlinear differential equations (Cambridge University Press, 1994)

44. Izhikevich, E. Dynamical Systems in Neuroscience: The Geometry of Excitability and Bursting (MIT Press, 2007)

45. Miller, K. D. Synaptic economics: competition and cooperation in synaptic plasticity. Neuron 17, 371–374, DOI: https://doi.org/10.1016/S0896-6273(00)80169-5 (1996)

46. Frankland, P. W. & Josselyn, S. A. Memory allocation. Neuropsychopharmacology 40, 243–243, DOI: http://dx.doi.org/10.1038/npp.2014.234(2015)

47. Erlenhardt, N. et al. Porcupine controls hippocampal ampar levels, composition, and synaptic transmission. Cell Rep 14, 782–794, DOI: http://dx.doi.org/10.1016/j.celrep.2015.12.078(2016)

48. Sjöström, P. J., Turrigiano, G. G. & Nelson, S. B. Rate, timing, and cooperativity jointly determine cortical synaptic plasticity. Neuron 32, 1149–1164, DOI: https://doi.org/10.1016/S0896-6273(01)00542-6 (2001)

49. Graupner, M. & Brunel, N. Mechanisms of induction and maintenance of spike-timing dependent plasticity in biophysical synapse models. Front Comput. Neurosci 4, 136, DOI: https://doi.org/10.3389/fncom.2010.00136 (2010)

50. Turrigiano, G. G. The dialectic of Hebb and homeostasis. Philos Trans R Soc Lond B Biol Sci 372, DOI: 10.1098/rstb.2016.0258 (2017)

51. Zenke, F. & Gerstner, W. Hebbian plasticity requires compensatory processes on multiple timescales. Philos Trans R Soc Lond B Biol Sci 372, DOI: 10.1098/rstb.2016.0259 (2017)

52. Kim, D., Paré, D. & Nair, S. S. Assignment of model amygdala neurons to the fear memory trace depends on competitive synaptic interactions. J. Neurosci. 33, 14354–14358, DOI: https://doi.org/10.1523/JNEUROSCI.2430-13.2013 (2013)

53. Kim, D., Paré, D. & Nair, S. S. Mechanisms contributing to the induction and storage of pavlovian fear memories in the lateral amygdala. Learn. Mem. 20, 421–430, DOI: 10.1101/lm.030262.113 (2013)

54. Cai, D. J. et al. A shared neural ensemble links distinct contextual memories encoded close in time. Nature 534, 115–118, DOI: https://doi.org/10.1038/nature17955 (2016)

55. Yokose, J. et al. Overlapping memory trace indispensable for linking, but not recalling, individual memories. Science 355, 398–403, DOI: 10.1126/science.aal2690 (2017)

56. Recanatesi, S., Katkov, M., Romani, S. & Tsodyks, M. Neural network model of memory retrieval. Front Comp Neurosci 9, 149, DOI: https://doi.org/10.3389/fncom.2015.00149 (2015)

